# Diagnosis and Prognosis of Alzheimer’s Disease Using Brain Morphometry and White Matter Connectomes

**DOI:** 10.1101/407601

**Authors:** Yun Wang, Chenxiao Xu, Ji-Hwan Park, Seonjoo Lee, Yaakov Stern, Shinjae Yoo, Jong Hun Kim, Hyoung Seop Kim, Jiook Cha, The Alzheimer’s Disease Neuroimaging Initiative

## Abstract

Accurate, reliable prediction of risk for Alzheimer’s disease (AD) is essential for early, disease-modifying therapeutics. Multimodal MRI, such as structural and diffusion MRI, is likely to contain complementary information of neurodegenerative processes in AD. Here we tested the utility of the multimodal MRI (T1-weighted structure and diffusion MRI), combined with high-throughput brain phenotyping—morphometry and structural connectomics—and machine learning, as a diagnostic tool for AD. We used, firstly, a clinical cohort at a dementia clinic (National Health Insurance Service-Ilsan Hospital [NHIS-IH]; N=211; 110 AD, 64 mild cognitive impairment [MCI], and 37 cognitively normal with subjective memory complaints [SMC]) to test the diagnostic models; and, secondly, Alzheimer’s Disease Neuroimaging Initiative (ADNI)-2 to test the generalizability. Our machine learning models trained on the morphometric and connectome estimates (number of features=34,646) showed optimal classification accuracy (AD/SMC: 97% accuracy, MCI/SMC: 83% accuracy; AD/MCI: 97% accuracy) in NHIS-IH cohort, outperforming a benchmark model (FLAIR-based white matter hyperintensity volumes). In ADNI-2 data, the combined connectome and morphometry model showed similar or superior accuracies (AD/HC: 96%; MCI/HC: 70%; AD/MCI: 75% accuracy) compared with the CSF biomarker model (t-tau, p-tau, and Amyloid β, and ratios). In predicting MCI to AD progression in a smaller cohort of ADNI-2 (n=60), the morphometry model showed similar performance with 69% accuracy compared with CSF biomarker model with 70% accuracy. Our comparison of classifiers trained on structural MRI, diffusion MRI, FLAIR, and CSF biomarkers show the promising utility of the white matter structural connectomes in classifying AD and MCI in addition to the widely used structural MRI-based morphometry, when combined with machine learning.

**Highlights:**

- We showed the utility of multimodal MRI, combining morphometry and white matter connectomes, to classify the diagnosis of AD and MCI using machine learning.
- In predicting the progression from MCI to AD, the morphometry model showed the best performance.
- Two independent clinical datasets were used in this study: one for model building, the other for generalizability testing.

## INTRODUCTION

There is an urgent, unmet need for clinically useful biomarkers of risk for Alzheimer’s disease (AD) based on non-invasive and affordable measures suited for routine examination of individuals with subthreshold symptoms. Studies have focused on brain MRI-derived markers. Cortical thinning and reduced hippocampal volumes based on structural MRI are known for markers for AD, but these structural estimates alone are insufficient for implementation at clinical settings because of insufficient accuracy and generalizability (Teipel et al., 2015).

It is conceptualized that biomarkers of Aβ deposition become abnormal early, and then markers of neuronal neurodegeneration or dysfunction show abnormality later in AD (Jack et al., 2010). These markers of neurodegeneration, rather than those of Aβ or Tau proteinopathy, appear directly related to cognitive symptoms (Jack et al., 2010). Neurobiology of AD relates to axonal and neuronal degeneration followed by fibrillar lesions triggered by amyloid precursor protein (APP)-initiated death-receptor mechanism and activation of tau (Holtzman et al., 2011; Nikolaev et al., 2009). Initial axonal degeneration may lead to grey matter tissue changes and finally to neuronal loss or atrophy resulting in cognitive and functional impairment. Since diffusion MRI uses water molecules as an endogenous tracer to probe tissue microstructure or properties (Beaulieu, 2002), it can detect subtle changes in microstructure tissue properties in AD. Previous studies have shown that decreased white matter integrity is associated with AD (Acosta-Cabronero et al., 2010; Douaud et al., 2011; Zhang et al., 2009).

A potentially powerful application of diffusion MRI to AD research is assessing axonal white matter tracts using tractography. Tractography is a computational reconstruction of white matter tracts using biophysical modeling of fiber orientations (Johansen-Berg and Behrens, 2006; Seehaus et al., 2013). Recent advances in computational methods have enabled more rigorous estimation of white matter tracts (Azadbakht et al., 2015; Ciccarelli et al., 2008; Shi and Toga, 2017; Sporns, 2011). In AD, human imaging of APP and tau shows widespread topography. Given this, when tractography is applied at the connectome level, this structural connectome data could be useful for assessing axonal or white matter abnormalities across the entire connectome. A few studies using tractography at the connectome level have noted abnormal topological organization of structural connectome in AD (Dai and He, 2014; Lo et al., 2010). However, it remains untested whether and to what extent the structural connectome carries additional information that structural MRI and morphometry analysis do not present.

In this study, we addressed this issue using rigorous, data-driven machine learning in two independent datasets of moderate sample sizes (211 elders for the first dataset [Korean National Health Insurance Service Ilsan Hospital, South Korea] and 179 elders for the second, generalizability dataset [ADNI-2]). In both data, using multi-modal brain MRI (structural and diffusion MRI), we performed high-throughput brain phenotyping, including automated morphometry and white matter structural connectomics (probabilistic tractography) to generate large-scale multi-modal, multi-parametric imaging-derived phenotypes used as features in machine learning. A well-established, rigorous analysis pipeline was applied to diffusion MRI to estimate robust, individualized structure connectomes. We compared data-driven machine learning classifiers trained on the individualized brain connectome and morphometric estimates with benchmark models (white matter hyperintensity) for the first Korean data and CSF biomarkers for the second reproducibility ADNI-2 data) using existing metrics.

## MATERIALS AND METHODS

### Participants

For the NHIS-IH Cohort, we used data from 211 seniors who visited the dementia clinic at National Health Insurance Service Ilsan Hospital (NHIS-IH), Goyang, South Korea from 2010 to 2015. This sample is a randomly selected subset of the Ilsan Dementia Cohort, a retrospective clinical cohort. Neurologists made a diagnosis based on possible AD and Peterson’s MCI criteria (Petersen, 2004), clinical history, a full battery of neuropsychological evaluations (Seoul neuropsychological screening battery) and MMSE (Mini-Mental State Examination). Those with vascular changes were not excluded from the study as long as they had a diagnosis of AD or MCI. Diagnosis is based on MMSE, CDR, and the neuropsychological evaluations. Distinction between MCI and SMC was based on the full battery of the neuropsychological evaluation (Seoul Neuropsychological Screening Battery-Dementia Version)(Ahn et al., 2010). To meet the diagnosis of MCI, an individual must show a neuropsychological score 1 SD below the normal range at least one of the nine domains of the full battery. Thus, all individuals with SMC show neuropsychological scores within the normal range; they are thus cognitively normal. Those with AD as a primary diagnosis and with small vessel disease were noted as “AD with small vessel disease”. Participants included 110 with the diagnosis of Alzheimer’s disease (AD; median age=82; interquartile intervals (Q3-Q1)=85-77), 64 with mild cognitive impairment (MCI; median age=73; Q3-Q1=77-66), and 37 subjective memory complaints (SMC; median age=74; Q3-Q1=78-72) (**Table 1**). To test the generalizability of our approach, we also used ADNI-2 (Alzheimer’s Disease Neuroimaging Initiative), where structural and diffusion MRI was collected. Demographical information is also provided in **Table 1**. The institutional review board of our hospital approved this study before implementation.

**Table 1.**
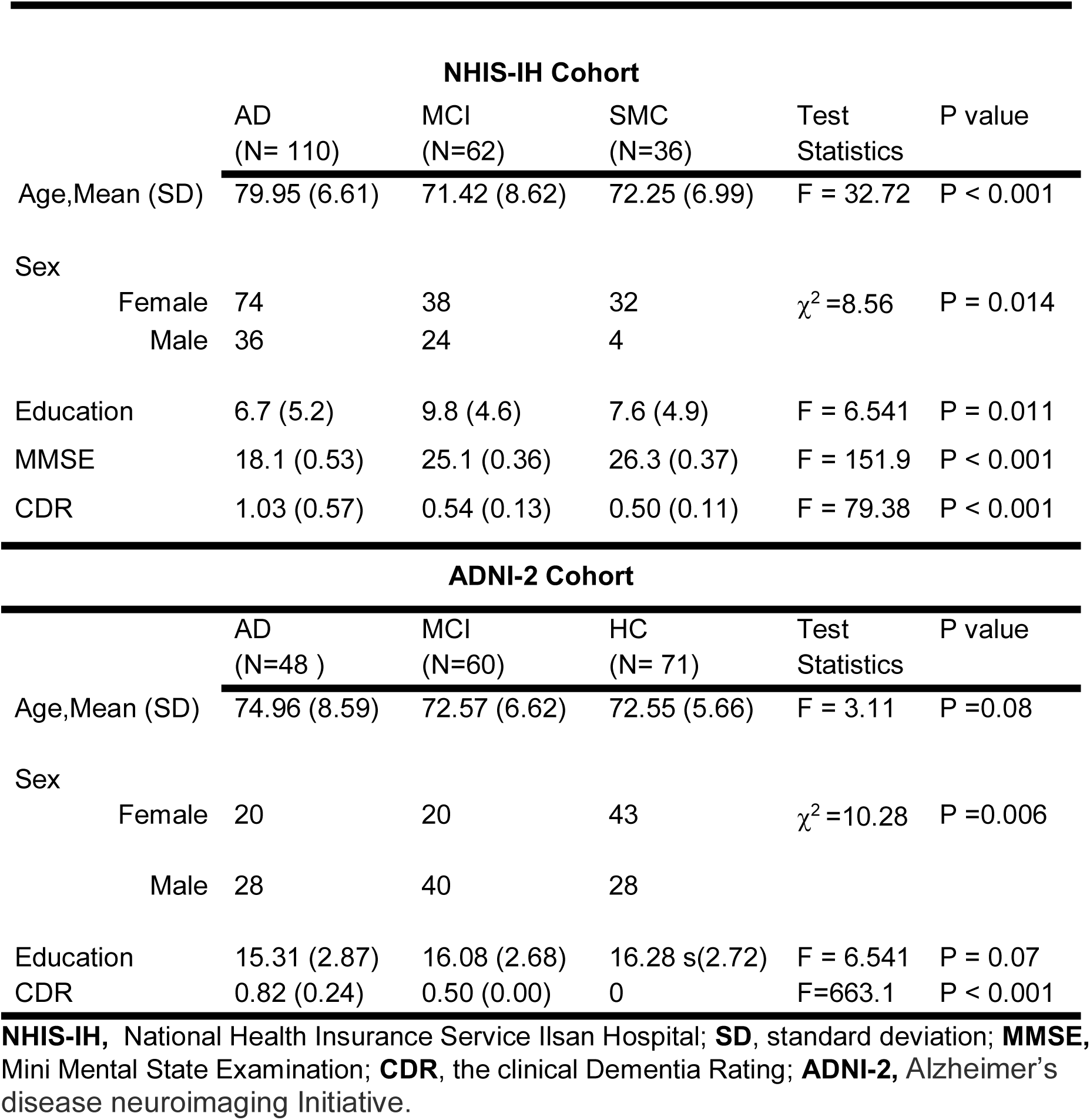
Participant Demographics

### MRI acquisition

National Health Insurance Service Ilsan Hospital (NHIS-IH): We collected the following multimodal MRI from all participants: T1-MPRAGE: TE, 4.6 ms; matrix, 310 ×480 ×480; voxel size, 0.5 ×0.5 ×0.5 mm. T2-FLAIR; matrix = 320 × 240 × 240; voxel size = 0.56 × 1.04 × 1.04. Diffusion MRI: matrix = 112 ×112 ×70; voxel size = 1.9 ×1.9 ×2.0 mm; the series included one image acquired without diffusion weighting and with diffusion weighting along 40 non-collinear directions (b = 600 s/m-2). ADNI-2: T1-weighted anatomical MRI and diffusion MRI. T1-MPRAGE: TE, min full echo; matrix, 208 ×240 ×256; voxel size, 1×1×1 mm. Diffusion MRI: matrix = 256 ×256×46; voxel size = 1.36×1.36 ×2.7 mm; the series included 5 image acquired without diffusion weighting and with diffusion weighting along 41 non-collinear directions (b = 1000 s/m-2).

### MRI Analysis-Structural MRI

The high-throughput computational analysis was conducted. First, we estimated morphometric estimates using the Freesurfer image analysis pipeline (Fischl, 2012) (v6) from T1 and T2-FLAIR images. Morphometric measures (N=948 per subject) include volumes of the hippocampal subdivisions, and thickness, surface area, and volume of cortical/subcortical regions using two different atlases available in Freesurfer (Desikan-Killiany atlas and Destrieux atlas; https://surfer.nmr.mgh.harvard.edu/fswiki/CorticalParcellation). The technical details of these procedures are described in previous studies (Desikan et al., 2006; Destrieux et al., 2010; Fischl and Dale, 2000; Fischl et al., 1999). In brief, the image processing includes motion correction, removal of non-brain tissue, Talairach transformation, segmentation, intensity normalization, tessellation of the gray matter-white matter boundary, topology correction, and surface deformation. Deformation procedures use both intensity and continuity information to produce representations of cortical thickness. The maps produced are not restricted to the voxel resolution and are thus capable of detecting submillimeter differences between groups.

### MRI Analysis-Diffusion MRI

We estimated structural connectome from structural and diffusion MRI. Structural MRI was used to define seed and target nodes of the connectome in each brain. We used the diffusion MRI analysis pipeline, MRtrix 3 (Tournier et al., 2004). The connectome measures (33,698 features per subject) include counts of streamlines, a surrogate measure of structural connectivity (Cha et al., 2015; Cha et al., 2017; Cha et al., 2016), and mean length of streamlines given any two brain regions based on multiple atlases. Diffusion-weighted magnetic resonance imaging (DWI) was preprocessed using the following pipeline in MRtrix 3. DWI was first denoised using a novel algorithm based on random matrix theory that permits data-driven, non-arbitrary threshold for Principal Component Analysis denoising; this method enhances the DWI quality for quantitative and statistical interpretation (Veraart et al., 2016). Denoised images then underwent eddy current and motion correction (Andersson and Sotiropoulos, 2016), brain extraction from three non-diffusion-weighted images (taking their median), and bias field correction using N4 algorithm (N4ITK), an improved N3 method, in Advanced Normalization Tools (ANTs)(Tustison et al., 2010). We then estimated fiber orientation distributions from each preprocessed image using 2^nd^-order integration over fiber orientation distributions (iFOD2). Based on the FODs, probabilistic tractography was performed using constrained spherical devolution (CSD). We used a target streamline count of 10 million across the whole brain. The tractograms were filtered using spherical-deconvolution informed filtering of tractograms (SIFT) with a target streamline count of 3 million. After a primary statistical analysis using these filtered tractograms, we tested whether the effects of interest were robust to the tractography and filtering parameters, such as the target streamline count for tractography, SIFT, or a ratio between them. This method permits mapping to streamline estimation back to individual’s DWI and updating a reconstruction to improve model fit. This approach renders the streamline counts connecting two brain regions proportional to the total cross-sectional area of the white matter fibers connecting those regions, enhancing streamline counts as a biologically plausible quantity, representing “structural connectivity”. This was done by repeating tractography and SIFT with a set of extreme parameters (100 million and 5 million target streamlines, respectively) with a filtering factor of 20 (100/5). Finally, from the filtered tractograms, we generated a connectivity matrix in each participant using brain parcellation and segmentation obtained from structural MRI from the same person. In this way, our structural connectome estimates reflect individualized connectomes. We used two different atlases in Freesurfer (Desikan-Killiany atlas (Desikan et al., 2006) and Destrieux atlas (Destrieux et al., 2010). We used streamline counts as the primary connectivity metric in this study as in a recent human infant imaging study (van den Heuvel et al., 2015b), as well mean length as secondary measures. A prior macaque study suggests the validity of streamline counts as an indicator of fiber connection strength, with the number of streamlines significantly correlating with tract-tracing strength in the macaque brain (van den Heuvel et al., 2015a).

### Machine Learning Classification

Given our goal to compare the classifiers trained on the distinct multimodal brain phenotypes, rather than to find a novel machine learning algorithm, we used the following three standard algorithms that have been extensively used in the literature(Abraham et al., 2014; Dimitriadis et al., 2018; Pellegrini et al., 2018): random forest, logistic regression (LR) with L1 and L2 regularization, and support vector machine (SVM) with a linear kernel. Also, given the majority of the prior machine learning classification studies in the AD literature are based on binary classification (Pellegrini et al., 2018), we chose binary classification for better comparison. Machine learning models were trained and cross-validated within each dataset. As a common preprocessing step for machine learning estimators, we standardized the imaging derived phenotypes by removing the median and scaling them according to the quantile range (i.e., between the 1^st^ and the 3^rd^ quartile); this method is known to be robust to outliers. Model training and validation were done using nested cross-validation to avoid overfitting due to bias to training data (Cawley and Talbot, 2010; Varoquaux et al., 2017). Nested cross-validation uses a series of train/validation/test set splits: In the inner loop, we trained the model and selected a set of hyperparameters using the training set, then optimized the model with validation set; In the outer loop, we estimated generalization error of the underlying model using test sets. For feature selection, we used the ‘forests of randomized trees’ method, an ensemble method to combine the predictions of base estimators built with a learning algorithm, and then tested whether additional PCA-based dimensionality reduction improved the model or not. For hyper-parameter optimization, we used the grid search method, varying C parameter for SVM and LR classifier, and varying the number of estimators and the minimum samples per leaf for random forest classifier. We used nested, k-fold, stratified cross-validation with ten iterations. To avoid information leakage during cross-validation, our nested cross-validation scheme used a series of train/validation/test set splits. First, in the inner loop, feature selection was performed, and the model was trained in a train set, and the model performance was maximized via hyper-parameter optimization in a validation set. Secondly, in the outer loop, the model performance was evaluated in a test set, and generalization error was estimated by averaging test set scores across cross-validation splits. To measure model performance, we used accuracy, sensitivity, specificity, F1 score, and Area Under the Curve in receiver operating characteristic (AUC ROC). In diagnostic classification, we tested six different binary classifications, AD (coded as 1) vs. SMC (coded as 0), AD vs. MCI, MCI vs. SMC, AD only vs. AD with small vessel diseases, AD only vs. MCI, AD only vs. SMC. All the ML analyses were done using scikit-learn, a python library for machine learning (Abraham et al., 2014).

### Benchmark models

We used existing biomarkers as benchmark models. First, white matter hyperintensity in the Korean NHIS-IH cohort, and CSF biomarkers in the ADNI-2 cohort. White matter hyperintensity measures were estimated from T2-weighted FLAIR images using Wisconsin White Matter Hyperintensities Segmentation Toolbox (Ithapu et al., 2014). This method uses supervised machine learning methods to segment hyperintense regions and generates normalized effective white matter hyperintensity volume. Second, in ADNI-2 data, we used CSF biomarkers (phosphorylated tau, total tau, AB, ratio of phosphorylated tau/AB, ratio of total tau/AB), whose utility as biomarkers for diagnosis of AD (Olsson et al., 2016), MCI, and progression to AD from MCI (Hansson et al., 2006) has been studied. Furthermore, CSF biomarkers are reported to precede symptom onset of MCI (Moghekar et al., 2013).

## RESULTS

### Classification of AD and MCI

In the NHIS-IH Cohort, we tested machine learning classification using white matter structural connectomes and morphometric estimates in 211 elders at the dementia clinic at the Korean National Health Insurance Service Ilsan Hospital. Age and sex alone showed moderate accuracies: AD/SMC: accuracy = 0.77; MCI/SMC: accuracy = 0.63; AD/MCI: accuracy = 0.72. White matter hyperintensity (WMH) served as a benchmark model, for it has been widely tested in the literature.

In classification of AD vs. SMC, optimal classification performance was shown in “morphometry+connectome” model (accuracy = 0.97, 95% CI=0.95-0.98) and “connectome” model (accuracy = 0.97, 95% CI=0.96-0.98) (**Table 2; Figure 1A**). These two models outperformed “morphometry” (accuracy = 0.87, 95% CI=0.85-0.88) and WMH benchmark models (accuracy = 0.73, 95% CI=0.71-0.75). In classification of MCI vs. SMC, similar classification performance was observed in “morphometry+connectome” (accuracy = 0.82, 95% CI=0.80-0.85) and “connectome” models (accuracy = 0.83, 95% CI=0.81-0.85), compared with lower performance of “morphometry” (accuracy = 0.59, 95% CI=0.57-0.60) and the WMH benchmark models (accuracy = 0.57, 95% CI=0.54-0.60). In classification of AD vs. MCI, “morphometry+connectome” models showed a best accuracy (accuracy=0.97, 95% CI=0.96-0.98), followed by “connectome” model (accuracy = 0.96, 95% CI=0.95-0.97), “morphometry” model (accuracy = 0.83, 95% CI=0.80-0.86), and the WMH benchmark models (accuracy = 0.66, 95% CI=0.64-0.69). Throughput all classifications, connectomes and morphometry showed greater diagnostic accuracies compared with the WMH benchmark.

**Table 2.**
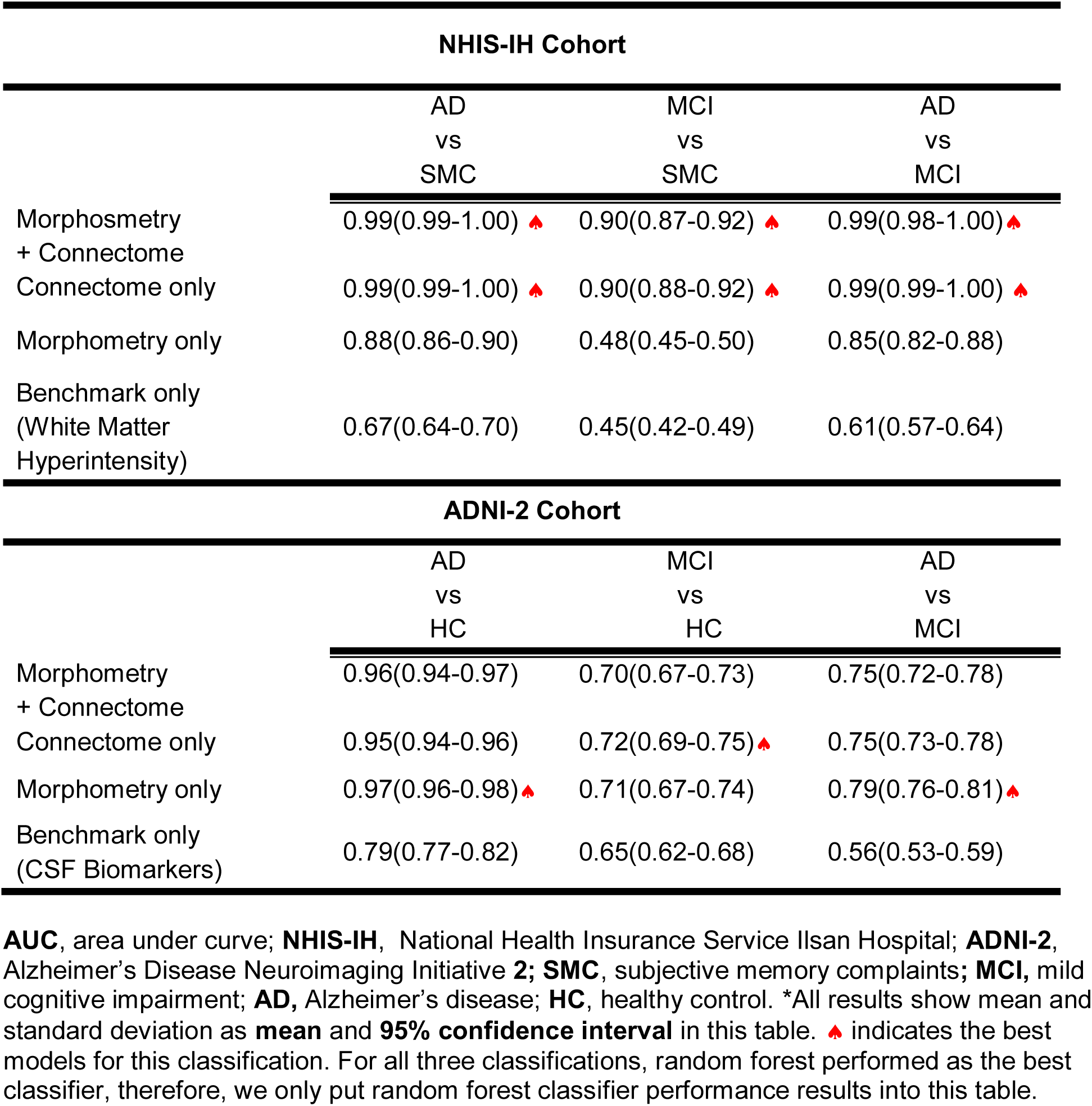
AUC Performances of Machine Learning Classifier using Structural Connectomes, Morphometric Brain Features, and benchmarks.

| **NHIS-IH Cohort** |  |  |  |
| --- | --- | --- | --- |
|  | AD<br>vs<br>SMC | MCI<br>vs<br>SMC | AD<br>vs<br>MCI |
| Morphosmetry<br>+ Connectome | 0.99(0.99-1.00) ▲ | 0.90(0.87-0.92) ▲ | 0.99(0.98-1.00) ▲ |
| Connectome only | 0.99(0.99-1.00) ▲ | 0.90(0.88-0.92) ▲ | 0.99(0.99-1.00) ▲ |
| Morphometry only | 0.88(0.86-0.90) | 0.48(0.45-0.50) | 0.85(0.82-0.88) |
| Benchmark only<br>(White Matter<br>Hyperintensity) | 0.67(0.64-0.70) | 0.45(0.42-0.49) | 0.61(0.57-0.64) |
| **ADNI-2 Cohort** |  |  |  |
|  | AD<br>vs<br>HC | MCI<br>vs<br>HC | AD<br>vs<br>MCI |
| Morphometry<br>+ Connectome | 0.96(0.94-0.97) | 0.70(0.67-0.73) | 0.75(0.72-0.78) |
| Connectome only | 0.95(0.94-0.96) | 0.72(0.69-0.75) ▲ | 0.75(0.73-0.78) |
| Morphometry only | 0.97(0.96-0.98) ▲ | 0.71(0.67-0.74) | 0.79(0.76-0.81) ▲ |
| Benchmark only<br>(CSF Biomarkers) | 0.79(0.77-0.82) | 0.65(0.62-0.68) | 0.56(0.53-0.59) |

**AUC**, area under curve; **NHIS-IH**, National Health Insurance Service Ilsan Hospital; **ADNI-2**, Alzheimer's Disease Neuroimaging Initiative 2; **SMC**, subjective memory complaints; **MCI**, mild cognitive impairment; **AD**, Alzheimer's disease; **HC**, healthy control. \*All results show mean and standard deviation as **mean** and **95% confidence interval** in this table. ▲ indicates the best models for this classification. For all three classifications, random forest performed as the best classifier, therefore, we only put random forest classifier performance results into this table.

**Table 3.** Performance in Predicting MCI to AD Progression in ADNI-2

| **Table 3. Performance in Predicting MCI to AD Progression in ADNI-2** |  |
| --- | --- |
| **MCI-AD vs. Stable MCI** |  |
| **Morphometry only** |  |
| (Best: LR + PCA+20 fold CV) |  |
| Accuracy | 0.69 (0.65-0.73)* |
| Sensitivity | 0.79 (0.74-0.83) |
| Specificity | 0.69 (0.64-0.74) |
| AUC | 0.79 (0.74-0.84) |
| **Connectomes only** |  |
| (Best: LR + PCA+20 fold CV) |  |
| Accuracy | 0.57 (0.53-0.61) |
| Sensitivity | 0.64 (0.58-0.69) |
| Specificity | 0.53 (0.47-0.59) |
| AUC | 0.62 (0.56-0.68) |
| **Morphometry + Connectome** |  |
| (Best: LR + PCA+10 fold CV) |  |
| Accuracy | 0.59 (0.56-0.62) |
| Sensitivity | 0.60 (0.56-0.63) |
| Specificity | 0.68 (0.56-0.79) |
| AUC | 0.65 (0.59-0.71) |
| **Benchmark: CSF biomarkers** |  |
| (Best: RF + no PCA+10 fold CV) |  |
| Accuracy | 0.70 (0.66-0.75) |
| Sensitivity | 0.76 (0.72-0.81) |
| Specificity | 0.71 (0.64-0.78) |
| AUC | 0.76 (0.70-0.81) |

**ADNI-2**, Alzheimer's Disease Neuroimaging Initiative 2; **MCI**, mild cognitive impairment; **AD**, Alzheimer's disease; **LR**, logistic regression; **PCA**, principal component analysis; **CV**, cross-validation. \*All results show Mean and standard deviation as **mean** and **95% confidence interval** in this table.

**Figure 1.**
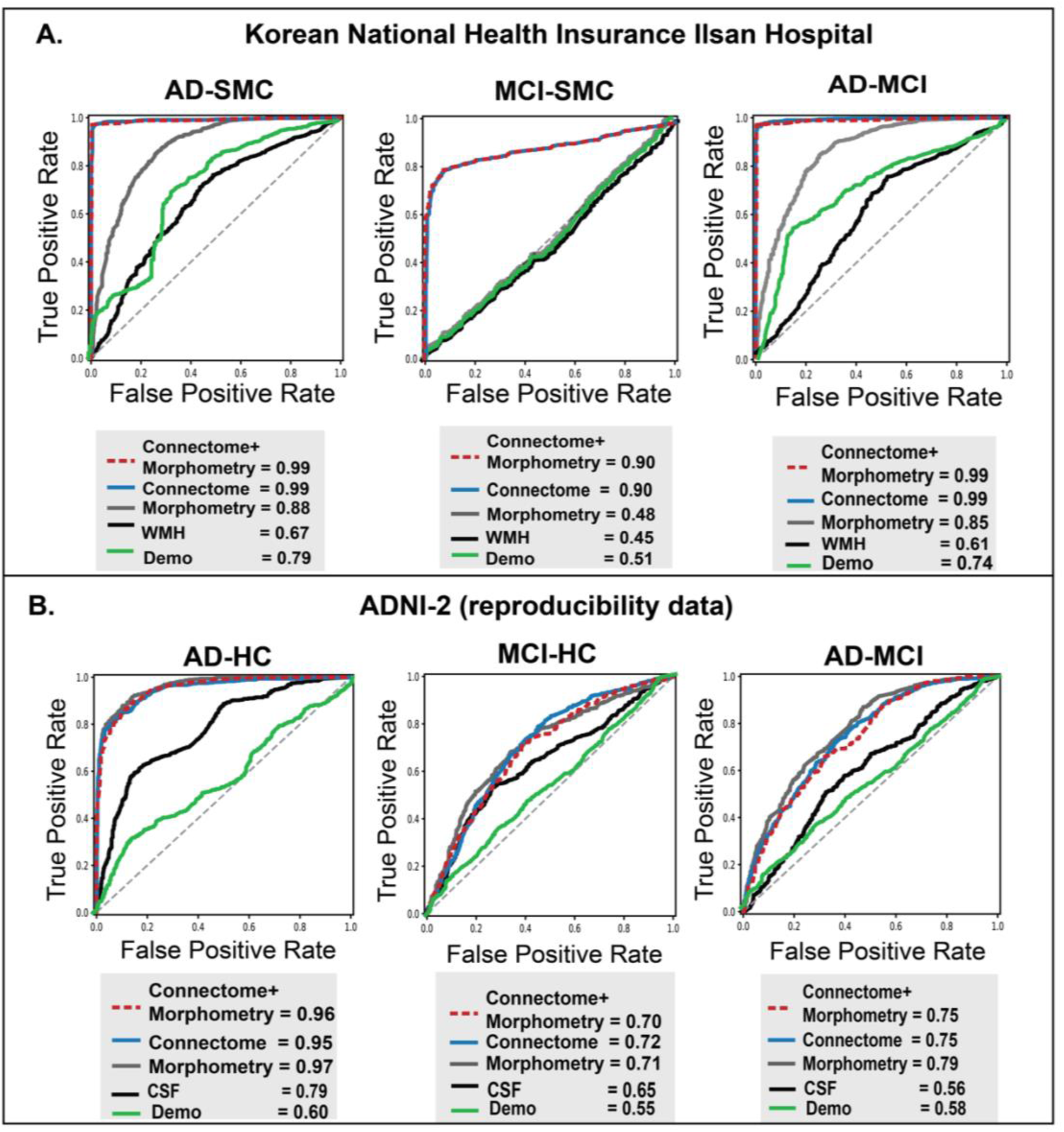
Classification of baseline diagnosis using connectomes and morphometric estimates. **Panel (*A)***, classification performances in the NHIS-IH Cohort (Korean National Health Insurance Ilsan Hospital data).It showed higher diagnostic accuracy (area under the curve of the receiver-operator characteristics or AUC ROC) of the machine learning model trained on combined connectome and morphometric estimates consistently, compared with the benchmark model trained on white matter hyperintensity. Out of three machine learning algorithms (random forest, support vector machine, and logistic regression), best models were shown. Panel (***B)***, classification performances in the ADNI-2 Cohort. It showed reproducible results of diagnostic accuracy of connectomes and morphometry. The combined models show better performance in predicting AD from healthy controls and AD from MCI, and similar in predicting MCI from HC. Best models were shown. Compared with the NHIS-IH Cohort, the reproducibility data shows less diagnostic accuracy presumably due to multiple sites and stricter inclusion and exclusion criteria in ADNI. **WMH**, white matter hyperintensity; **Demo**, demographics including sex, age, and education.

### Testing generalizability

We next tested the generalizability of the same multimodal brain imaging-based machine learning using ADNI-2 data. We included participants in ADNI-2 data whose structural and diffusion MRI (baseline) were both collected. To compare the performance of our classifiers, we used the invasive CSF biomarkers (p-tau, t-tau, Aβ42, p-tau/ Aβ42, t-tau/ Aβ42) as a benchmark model. In the classification of AD vs. HC, all the MRI-based models showed similarly optimal performance around 0.88 accuracy (**Table 2; Figure 1B**), outperforming the CSF benchmark model (accuracy = 0.75, 95% CI=0.73-0.77). In classification MCI vs. HC, all the MRI-based models showed similar performance with accuracies ranging from 0.64-0.67, outperforming the CSF benchmark (accuracy = 0.62, 95% CI=0.59-0.65). In classification AD vs. MCI, all the MRI-based models showed similar performance with accuracy ranging from 0.66-0.71, outperforming the CSF benchmark (accuracy = 0.54, 95% CI=0.52-0.57) which is barely above chance. This generalizability data showed, firstly, morphometry and connectome estimates showed equally good performance consistently exceeding the invasive CSF biomarkers in classifying AD/MCI/HC; secondly, unlike the NHIS-IH results, synergistic effects of combined morphometry and connectomes were not observed using our machine learning framework.

### Testing utility for prognosis

Of the ADNI-2 data, we further tested the utility of our approach in predicting the disease trajectory. Data from 60 elders were used, whose baseline diagnosis was MCI and who were followed for at least two years. Machine learning models trained on the same five CSF benchmarks were used as a benchmark. In predicting progression from MCI to AD, “morphometry” model showed a highest accuracy (accuracy = 0.69, 95% CI=0.65-0.73) among MRI-based models, similar to the CSF benchmark model (accuracy = 0.70, 95% CI=0.66-0.75). (**Table 5, Figure 2**). “Connectome” model showed a lower, but statistically meaningful accuracy (accuracy = 0.57, 95% CI=0.53-0.61). Combining the two modalities of morphometry and connectomes (“morphometry+connectome”) did not improve the prognosis accuracy (accuracy = 0.59, 95% CI=0.56-0.62), compared with “morphometry” model.

**Figure 2.**
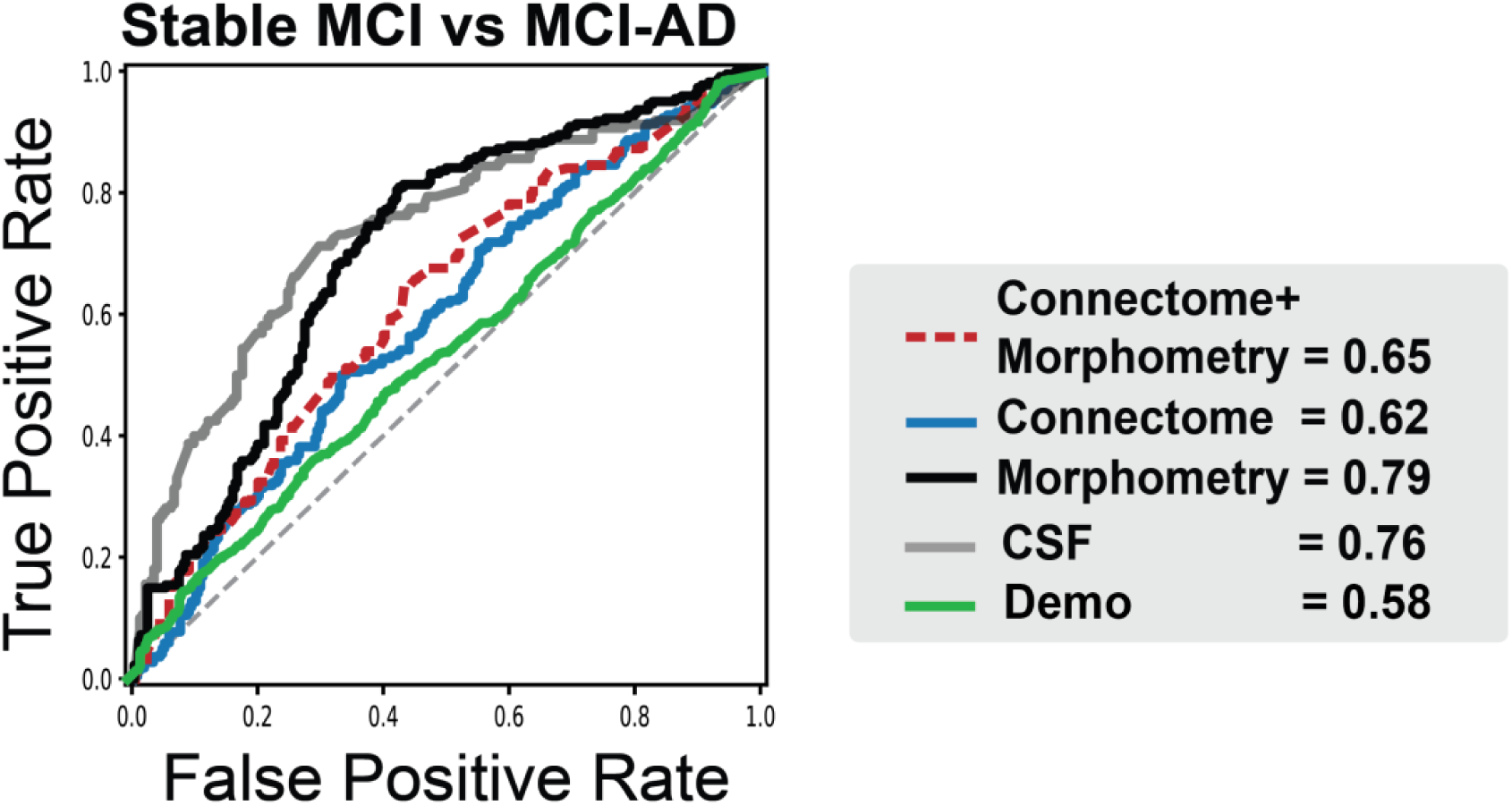
Prediction of progression to AD from MCI using connectomes and morphometric estimates. Using ADNI-2 data that has follow-up data after baseline MRI scan, machine learning models were tested using connectome and morphometry estimates to predict MRI-to-AD progression in 60 elders with MCI (mean follow-up years in stable MCI, 3.76 ± 0.98; range, 2.18-5.32). Morphometry model showed similar performance to CSF benchmark model. Both the combined model and connectome model showed lower but meaningful accuracy.

## DISCUSSION

In this study, we used large-scale MRI-derived brain phenotypes (morphometry and white matter structural connectomes) with machine learning techniques to test AD and MCI diagnosis in two independent Alzheimer’s disease datasets. We also predicted disease progression to AD from MCI. For high-throughput imaging analysis, we used a well-established automated pipeline for morphometry and a pipeline to estimate rigorously individualized white matter structural connectomes. Firstly, the models trained on morphometry and connectomes showed the best accuracy in classifying AD, MCI, and SMC or HC in the single-site data (ranging from 90% to 99% in AUC ROC; NHIS-IH, South Korea) as well as the multi-site (ranging from 70% to 97% in AUC ROC; ADNI-2, USA) “reproducibility” data. The models outperformed the benchmark models significantly (e.g., white matter hyperintensity or CSF biomarkers) and demographic model (including age, sex, and education). Second, the model trained on connectome or morphometric estimates showed moderate accuracies (ranging from 57% to 79%; AUC) in predicting progression to AD in 60 elders with MCI in ADNI-2 data. These results show the utility of white matter structural connectomes in addition to morphometry in detecting the abnormal brain aging process in AD pathology.

A novel aspect of this study is to assess the utility of the dMRI-based white matter structural connectomes in predictive modeling of AD in a sufficiently large sample (n=211) and to validate it in an independent cohort (n=179). In the NHIS-IH data, the “connectome” model and “connectome and morphometry” model similarly show the optimal classification of AD or MCI, outperforming the benchmark model of white matter hyperintensity. Likewise, in the ADNI-2 generalizability data, both “connectome” and “connectome and morphometry” models show optimal classification accuracy, outperforming the CSF benchmark model. This finding is in line with the literature showing the associations of structural connectomes with potential AD pathology (e.g., topological disturbance based on graph theory) (Pereira et al., 2017) and with healthy aging (Perry et al., 2015). Also, prior studies show the potential utility of connectomics estimates in predicting risk for AD, but with a caveat of limited samples sizes (n<30 (Wee et al., 2012; Zhu et al., 2014)). Our study thus further demonstrate the potential practical utility and generalizability of the unbiased brain analytic approach combined with data-driven machine learning, leveraging two independent data with greater sample sizes.

The classification results in the NHIS-IH data may further suggest an important implication. The morphometry model fails to classify MCI from SMC, whereas the connectome or combined model shows optimal classification of 0.90 AUC. The gain of the connectome estimates in classification is more pronounced in MCI/SMC classification than in AD/SMC classification. This might suggest a greater sensitivity of the white matter connectivity estimates in detecting AD-related neurodegeneration compared with grey matter morphometry. Literature shows the capability of diffusion MRI-derived measures to detect subtle microscopic changes in tissue properties or integrity (Acosta-Cabronero et al., 2010; Beaulieu, 2002; Douaud et al., 2011; Zhang et al., 2009), whereas structural MRI is typically used to estimate macroscopic properties, namely volumes. However, this pattern is not seen in the ADNI-2 multi-site data; this leads to an issue of data harmonization to deal with site effects of MRI-derived estimates.

The connectome or combined model shows ∼10% decrease in model performance in the ADNI-2 multi-site data compared with the NHIS-IH single-site data. It is possible that it is related to the site variability in the dMRI data. Indeed, prior studies show persistent inter-site variability in diffusion data even when using similar types of scanners, pulse sequences or same field strength (Fox et al., 2012; Mirzaalian et al., 2016). This is a non-trivial problem because there are hardly any objective ways to assess harmonization of dMRI data (e.g., a dynamic phantom optimized for dMRI). One potential way to mitigate this variability issue across multiple data sources is an analytical solution. A recent study suggests an elegant Bayesian method for post-acquisition harmonization of dMRI (Fortin et al., 2017). In our study, however, this method could not be applied to our raw dMRI or fiber orientation distribution maps for probabilistic tractography.

One potential approach to MRI harmonization is domain-invariant machine learning. A recent seminal study (Ghafoorian et al., 2017) of white matter hyperintensity segmentation in the brain shows a successful application of “multi-source domain adaption”. That is, a convolutional neural network trained on data from a single domain (i.e., from a single scanner with a single acquisition protocol) was successfully applied (retrained) to the same task with independent MRI from different domains (i.e., different acquisition protocols and image dimension from the same scanner). Given the recent rapid development of the deep learning algorithms, Artificial Intelligence-based domain adaptation might be a promising way towards the generalizable and reproducible MRI-based analytics.

In predicting MCI-to-AD progression in the ADNI-2 data, the morphometry model outperforms both connectome and combined models. This may first suggest that grey matter morphometry provides more useful information in predicting the AD trajectory than the connectome measures. However, given the smaller sample size (N=60) compared with AD/MCI classification (N=119), in this analysis we suspect that machine learning training and feature selection may be suboptimal for the connectome model than for the morphometry model, because of the significantly large number of features in the former (N=33,698) than the latter (N=948). Similarly, while the morphometry model and connectome model respectively showed statistically meaningful (above chance) predictions, when combined, there was little improvement in model performance. This indicates more rigorous methods to combine models trained across multimodal brain imaging-derived phenotypes may be required, such as ensemble methods (Zhang et al., 2011).

Limitations related to the NHIS-IH data include the significantly greater age in the AD group compared with the MCI or SMC groups. It is possible that a greater aging effect embedded on the brain phenotypes may have made the classification of AD easier. However, in ADNI data with the age-matched samples, classification performance (AUC=0.97) was only slightly less than the NHIS-IH data (AUC=0.99). This suggests that the patterns extracted from morphometry and white matter connectomes may be specific to AD rather than an age-related bias. Another limitation is the lack of healthy controls in the NHIS-IH cohorts. In this retrospective cohort at the dementia clinic, individuals with Subjective Memory Complaints are cognitively normal.

Nevertheless, this group might not be equivalent to healthy controls as in the ADNI data. For example, there might be subtle differences in brain health status between health individuals and cognitively normal individuals with subjective memory complaints. Our study provides no data to address this. Nevertheless, given the fact that in clinical settings, individuals seek for clinical service usually when they suspect symptoms, our results of classifying AD and MCI from individuals with SMC may have a unique clinical utility in addition to the comparisons of AD and MCI with healthy controls in the ADNI data.

In sum, this study lends support for the individualized white matter structural connectomes, estimated from multimodal MRI (structural and diffusion), in combination with machine learning techniques, as a useful method to detect accurately AD-related neurodegeneration across the whole brain in a data-driven manner.

## Acknowledgments

This work used the Extreme Science and Engineering Discovery Environment Stampede 2 at the Texas Advanced Computing Center (TG-IBN170015: Cha) and Argonne National Laboratory Leadership Computing Facility (PI, Cha). This study was supported by NIMH K01 MH109836 (Cha), Brain and Behavior Research Foundation NARSAD Young Investigator award (Cha), Korean Scientists and Engineers Association Young Investigator Grant (Cha), National health insurance Ilsan hospital research fund.

